# CoV-UniBind: A Unified Antibody Binding Database for SARS-CoV-2

**DOI:** 10.1101/2025.07.17.665346

**Authors:** Aryan Bhasin, Francesco Saccon, Callum Canavan, Andrew Robson, Joao Euko, Alexandra C. Walls, Yunguan Fu

## Abstract

Since the emergence of SARS-CoV-2, numerous studies have investigated antibody interactions with viral variants *in vitro*, and several datasets have been curated to compile available protein structures and experimental measurements. However, existing data remain fragmented, limiting their utility for the development and validation of machine learning models for antibody–antigen interaction prediction. Here, we present CoV-UniBind, a unified database comprising over 75,000 entries of SARS-CoV-2 antibody–antigen sequence, binding, and structural data, integrated and standardised from three public sources and multiple peer-reviewed publications. To demonstrate its utility, we benchmarked multiple protein folding and inverse folding models across tasks relevant to antibody design and vaccine development. We expect CoV-UniBind to facilitate future computational efforts in antibody and vaccine development against SARS-CoV-2.

**Availability and implementation:** The curated datasets, structures, model scores and antibody synonyms are free to download at https://huggingface.co/datasets/InstaDeepAI/cov-unibind. Folded structures are available upon request.

## Introduction

Since the emergence of the Severe Acute Respiratory Syndrome Coronavirus 2 (SARS-CoV-2), extensive global efforts have focused on the development of vaccines (1, 2) and monoclonal antibodies (3) to combat the COVID-19 pandemic. These initiatives have generated a vast amount of publicly available genomic (4), structural (5), and biochemical data (6–8), providing a valuable foundation for data-driven research. Simultaneously, rapid advancements in artificial intelligence (AI) have enabled the application of deep learning models across multiple areas of vaccine and therapeutic development (9), including variant monitoring (10), antibody binding prediction (11), epitope identification (12), and mutational impact on immune recognition (13). Despite the increased use of AI in SARS-CoV-2 research, systematic evaluation of the models used remains limited. As COVID-19 transitions into a long-term global challenge, there is a growing need for sustained and rigorous assessment of such AI methods. A comprehensive benchmarking database is therefore essential to evaluate their utility and performance in COVID-19–related applications over time.

One key challenge in effective model benchmarking is the fragmented landscape of existing datasets, marked by inconsistent annotations and limited structural data (only ∼ 1, 270 relevant structures are currently available in the RCSB Protein Data Bank (PDB) (14)). This presents complications when evaluating protein folding and inverse folding models, which require large amounts of high-quality, annotated structural data. While some databases aim to integrate multiple data sources and standardise benchmarking (15–17), they often suffer from limited experimental coverage, incomplete structural data, or lack of open access. SARS-CoV-2-specific benchmarking resources are particularly scarce. Notably, this challenge is not limited to AI model evaluation but also affects experimental studies, where the lack of standardised benchmarks hampers reproducibility and cross-study comparisons.

To address these limitations, we present CoV-UniBind, a unified antibody binding database for SARS-CoV-2, which integrates high-resolution antibody–antigen complex structures with diverse experimental data capturing complementary aspects of antibody–antigen interactions. These data aim to provide a foundation for understanding SARS-CoV-2 evolution and guiding the design of next-generation vaccines and therapeutic antibodies against emerging variants. In CoV-UniBind, binding annotations from the Coronavirus Antibody Database (CoV-AbDab) (5) provide curated evidence of lineage-specific antibody responses. Precise kinetic measurements of binding strength and interaction dynamics are provided by protein–protein interaction assays, such as surface plasmon resonance (SPR) experiments (18). Complementing these data, we include fine-grained, residue-level analyses which assess the impact of individual mutations on antibody binding, from a high-throughput strategy known as deep mutational scanning (DMS) (19). In addition to the binding-focused datasets, the Drug Resistance Database (DRDB) (20) provides quantitative neutralisation potency measurements, offering an extended perspective on antibody efficacy against diverse viral variants. Available structural information in the Structural Antibody Database (SAbDab) (21) can be further processed to derive epitope and paratope information, antibody variable heavy/light chain (VH/VL) sequences and complementarity-determining regions (CDRs). By integrating these diverse yet interlinked data types, CoV-UniBind provides a comprehensive resource for benchmarking and advancing computational models of SARS-CoV-2 antibody–antigen interactions.

To demonstrate the utility of the curated database, we bench-marked a selection of folding (22, 23) and inverse folding models (24–26) on antibody–antigen binding ranking across two categories: ranking multiple antigen variants for each antibody and ranking different antibodies for each antigen variant. These tasks reflect critical applications in vaccine development and therapeutic antibody design, respectively. As new SARS-CoV-2 variants continue to emerge (as of June 2025), such as the LP.8.1 lineage (27) or more recently NB.1.8.1 (28), designated by the World Health Organisation as variants under monitoring on 24 January 2025 and 23 May 2025, respectively, we anticipate that CoV-UniBind will serve as a valuable resource for benchmarking predictive models to support next-generation vaccine and therapeutic antibody development in response to viral evolution (29).

## Materials and methods

### Data sources

The CoV-UniBind database was curated from multiple public sources, including SAbDab (21) for protein structures, CoV-AbDab (5) comprising lineage-specific binding and neutralisation labels, and the COVID DRDB (20) containing half-maximal inhibitory concentration (IC_50_) from virus neutralisation assays against monoclonal antibodies. Mutational data for multiple SARS-CoV-2 lineages was obtained from outbreak.info (30). In addition, data from several peer-reviewed studies were collected and integrated. Namely, Greaney et al. (19), Cao et al. (31) provided binding escape data from DMS experiments, (18), (32), (33) provided kinetic data from SPR assays, and (34) provided binding data (IC_50_) from enzyme-linked immunosorbent assay (ELISA) experiments.

### Antibody–antigen structure curation

The antibody– antigen protein structures were retrieved from SAbDab in the Chothia scheme (35), and restricted to entries for which the antigen is the SARS-CoV-2 spike glycoprotein. Water molecules and heteroatoms were removed. Amino acid sequences were additionally obtained from the PDB (36), used to extract antibody names, VH and VL chain sequences, and CDR amino acid sequences using ANARCI (37). The CDR residue ranges used are defined by (38). Given that a single PDB structure can contain a full spike trimer with multiple associated antibodies, each complex structure was split into individual antibody–antigen pairs (based on those in contact), with the antibody trimmed to include only the VH and VL domains, and the antigen trimmed to the receptor-binding domain (RBD) or N-terminal domain (NTD). Epitope and paratope residues were identified as contact residues between the CDRs and the antigen. Following Jiang et al. (39), two residues were defined as contacting if the minimum distance between any of their atoms was ≤ 5Å.

### Antibody identification

Antibodies often have synonyms, and annotations can vary across datasets. In CoV-UniBind, each antibody is uniquely identified by its six CDR amino acid sequences: CDR-H1, CDR-H2, CDR-H3, CDR-L1, CDR-L2, and CDR-L3. Antibodies with different VH and VL sequences but identical CDRs are considered function-ally equivalent with respect to antigen interaction. For CoV-AbDab, the antibody matching with their associated structural data was performed by comparing CDRs since CoV-AbDab provided VH and VL sequences. This provided an antibody synonym library that was used for all other datasets. For the DMS, SPR and IC_50_ datasets, standardised antibody synonyms and PDB IDs, when available, were used to match the entries with their respective structural information.

### Antibody–antigen interaction prediction

AlphaFold2 (AF2) was evaluated in two different modes: multimer model and monomer model with gap trick (42). The predicted local distance difference test (plDDT), predicted aligned error (PAE), and predicted template modelling (pTM) metrics were extracted from the predictions, where larger plDDT and pTM values, and lower PAE values, are intuitively hypothesized to correlate with stronger binding affinity. Both configurations were run with default parameters, without multiple sequence alignment (MSA), and using a custom structural template, a complex of the antibody with a reference antigen. For each antibody, a single custom template was reused across all associated variants. The `model_2_ptm` checkpoint was used for all predictions. Boltz-1 was evaluated similarly, with predicted docking error (PDE) instead of PAE in addition to the other metrics (43). It was executed with its default settings, also without the use of templates or MSA. All folding experiments (using AF2-Monomer with gap trick, AF2-Multimer and Boltz-1) were run with an internal folding pipeline, which enables fast and high-throughput protein folding predictions via the Google Cloud Platform (GCP) environment.

For inverse folding models (protein message passing neural networks (MPNN) (44), MPNNsol (25), AbMPNN (26), and FAMPNN (45)), the binding is estimated using the log-likelihood of the introduced antigen mutations with respect to a reference antigen sequence. Scoring was conditioned on the structure of an antigen–antibody complex, typically using the wild-type antigen as the structural reference. For FAMPNN, 10 sampling runs were performed with different random seeds, all with a batch size of 16, and the final score of each antigen was computed as the mean across these 10 samples. Three model checkpoints were tested, namely `0_0`, `0_3` and `0_3_cath` as described in (45). For other inverse folding models, 100 sequence samples were generated per input, with the final score defined as the mean across those 100 samples. A batch size of 100 and a fixed random seed of 42 were used.

As additional baselines, we included two non-deep learning methods. The first is epitope alteration count (10), which calculates the number of antigen mutations occurring within the structure-derived epitope. A higher count is expected to correspond to reduced binding, reflecting the assumption that more alterations in the binding interface lead to binding abrogation. This approach does not consider the biochemical nature or positional impact of the mutations. The second baseline leverages DMS data. For each single-point mutation, the escape value was first averaged across all antibodies in the corresponding DMS dataset. These values were then summed to produce a final score for multi-point mutants, with higher scores indicating greater escape potential and thus lower expected binding. This procedure was applied separately using the Bloom and Cao DMS datasets, resulting in two baselines: Bloom-based and Cao-based DMS scores.

### Statistical analysis

For antigen ranking tasks, we evaluate model performance using the values of receiver operating characteristic area under the curve (ROC-AUC), averaged across multiple antibody scores within the CoV-AbDab dataset, which is based on binary binding labels. For all other datasets, based on continuous binding values, we report the mean Spearman correlation coefficient 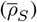, along with the standard error of the mean (SEM), computed across antibodies: 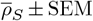. For antibody ranking tasks, we follow the same evaluation procedure, but aggregate performance across multiple antigens. Specifically, we report ROC-AUC for CoV-AbDab and 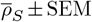 for all other datasets.

For each model, a single performance metric was selected to calculate correlations with target values. All metrics were aligned such that higher values consistently indicate better performance, enabling consistent comparisons of ROC-AUC values and Spearman correlation coefficients. Given that the continuous target values were either IC_50_, antibody escape or *K*_*D*_, lower values indicate better binding. For Boltz-1, the plDDT of the full antigen–antibody complex was used with reversed sign, since higher plDDT values reflect higher prediction confidence. For FAMPNN, the negative likelihood ratio was used, and for the backbone-only inverse folding models (ProteinMPNN, MPNNsol, and AbMPNN) the negative log-likelihood scores were used. AlphaFold-2 models (both monomer and multimer) relied on the overall complex PAE scores. Hence, a lower score is “better” for all of the deep learning model metrics used, in accordance with the binding target values. A two-tailed Mann-Whitney U-test was used to compare the ROC-AUC values of groups of antibodies binding to two different domains of the spike protein.

## Results and Discussion

The resulting CoV-UniBind database comprises 308 antibody–antigen complex structures, 73 unique antigen lineages (2403 antigen variants, including 2179 single-point mutants) and 175 unique antibodies. The vast majority of the antibodies represented in the datasets bind to the RBD. Only 7 antibodies bind to the NTD, of which 6 are present in the CoV-AbDab dataset. In particular, the DMS, Jian ELISA, and SPR datasets are limited to RBD-binding antibodies. An expanded summary of the composition and characteristics of each data source is presented in Table 1.

**Table 1.**
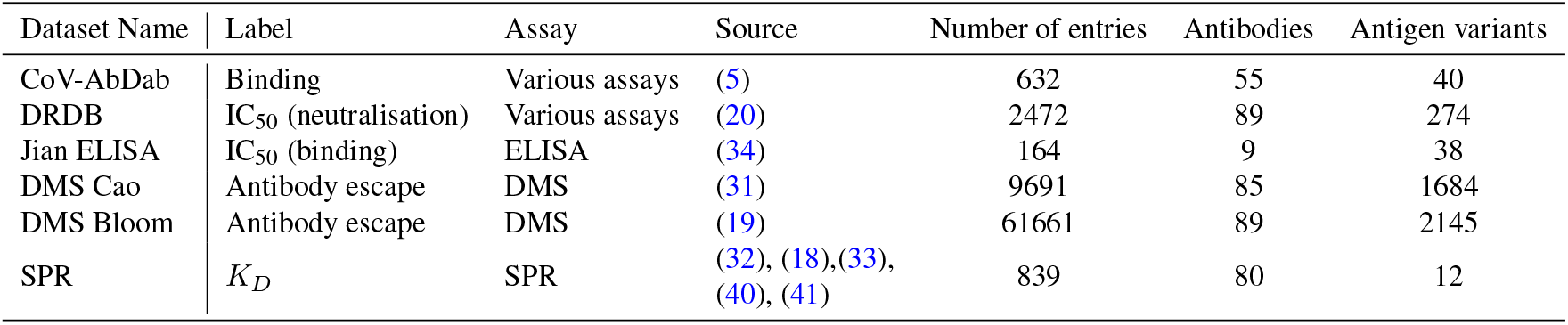
Summary of antibody–antigen interaction datasets. Unique antigen variants include both lineages from outbreak.info and single-point mutants generated in DMS studies.

| Dataset Name | Label | Assay | Source | Number of entries | Antibodies | Antigen variants |
| --- | --- | --- | --- | --- | --- | --- |
| CoV-AbDab | Binding | Various assays | (5) | 632 | 55 | 40 |
| DRDB | IC<sub>50</sub> (neutralisation) | Various assays | (20) | 2472 | 89 | 274 |
| Jian ELISA | IC<sub>50</sub> (binding) | ELISA | (34) | 164 | 9 | 38 |
| DMS Cao | Antibody escape | DMS | (31) | 9691 | 85 | 1684 |
| DMS Bloom | Antibody escape | DMS | (19) | 61661 | 89 | 2145 |
| SPR | $K_D$ | SPR | (32), (18), (33),<br>(40), (41) | 839 | 80 | 12 |

To benchmark models, we framed CoV-UniBind as two predictive tasks: (i) ranking antigens by predicted binding affinity for each antibody, and (ii) ranking antibodies for each antigen. This formulation addresses dataset limitations (e.g., DMS data are normalised per antibody) and supports distinct applications: antigen ranking aids vaccine design by assessing escape potential, while antibody ranking supports therapeutic discovery by identifying potent binders.

CoV-AbDab provides binary binding labels for a diverse set of antibodies, including 49 targeting the RBD and 6 targeting the NTD. The nature of this dataset makes it well-suited for antigen (or antibody) classification tasks. The diversity of antibody binding sites in CoV-AbDab also allows for benchmarking antigen ranking models across both RBD and NTD subsets (Table S1). In the antigen ranking task, all models demonstrate moderate to good performance (ROC-AUC ≥ 0.555) on the CoV-AbDab dataset (Table 2), where the AF2-monomer model with gap trick achieves the highest mean ROC-AUC of 0.850 ± 0.031. In contrast, all tested deep learning models struggled to generalise to the DMS datasets, with mean Spearman correlations 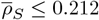. This underscores a key challenge posed by the DMS data, which consist of single-point mutations, where escape is measured indirectly as a proxy for binding affinity (46). The Jian ELISA and SPR datasets, like CoV-AbDab, involve comparisons across antibodies and viral lineages (including multiple mutations) and offer more direct measures of binding strength, either via potency (IC_50_) values or kinetic (k_D_) data. Inverse folding models, particularly FAMPNN, exhibit strong correlations on these datasets, achieving a mean Spearman correlation of 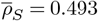 on the Jian ELISA dataset.

**Table 2.** Performance metrics for antigen ranking across binding datasets and models. Reported values include the mean ± SEM for both ROC-AUC and Spearman correlation coefficient 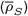. Higher positive values indicate stronger performance, where the best-performing model is highlighted in bold and second best is underlined. – indicates data are not available, due to computational constraints, AF2-multimer and Boltz-1 were not evaluated on the DMS Cao dataset, while AF2-multimer, AF2-monomer with the gap trick, and Boltz-1 were not run on the DMS Bloom dataset.

| Model | Cov-AbDab<br>Binding ROC-AUC | DMS Cao<br>Antibody Escape $\bar{\rho}_S$ | DMS Bloom<br>Antibody Escape $\bar{\rho}_S$ | Jian Binding<br>Potency $\bar{\rho}_S$ | SPR Binding<br>Affinity $\bar{\rho}_S$ |
| --- | --- | --- | --- | --- | --- |
| Epitope Alteration Count | 0.802 ± 0.037 | 0.242 ± 0.021 | 0.228 ± 0.020 | 0.418 ± 0.100 | **0.637 ± 0.032** |
| Cao DMS Baseline | 0.799 ± 0.041 | **0.483 ± 0.017** | <u>0.409 ± 0.020</u> | 0.459 ± 0.082 | 0.590 ± 0.031 |
| Bloom DMS Baseline | <u>0.803 ± 0.042</u> | <u>0.417 ± 0.021</u> | **0.444 ± 0.023** | <u>0.490 ± 0.084</u> | <u>0.598 ± 0.029</u> |
| AF2 Multimer | 0.555 ± 0.039 | – | – | 0.134 ± 0.110 | 0.088 ± 0.037 |
| AF2 Monomer | **0.850 ± 0.031** | 0.212 ± 0.020 | – | 0.319 ± 0.108 | 0.485 ± 0.038 |
| Boltz1 | 0.681 ± 0.039 | – | – | 0.059 ± 0.069 | 0.284 ± 0.040 |
| ProteinMPNN | 0.726 ± 0.038 | 0.064 ± 0.020 | 0.106 ± 0.016 | -0.017 ± 0.143 | 0.244 ± 0.039 |
| MPNNsol | 0.696 ± 0.036 | 0.073 ± 0.019 | 0.113 ± 0.014 | 0.054 ± 0.121 | 0.215 ± 0.039 |
| AbMPNN | 0.765 ± 0.037 | 0.036 ± 0.019 | 0.066 ± 0.014 | 0.160 ± 0.125 | 0.344 ± 0.038 |
| FAMPNN 0_3_cath | 0.749 ± 0.038 | 0.102 ± 0.023 | 0.135 ± 0.018 | 0.044 ± 0.126 | 0.326 ± 0.044 |
| FAMPNN 0_3 | 0.666 ± 0.049 | 0.003 ± 0.017 | 0.023 ± 0.015 | **0.493 ± 0.084** | 0.320 ± 0.054 |
| FAMPNN 0_0 | 0.738 ± 0.043 | -0.052 ± 0.020 | 0.005 ± 0.016 | 0.349 ± 0.134 | 0.431 ± 0.039 |

Table 3 summarises benchmarking results for the antibody ranking task. Among structure-based folding models, Boltz-1 achieved the highest classification performance on the CoVAbDab dataset, with an ROC-AUC of 0.562 ± 0.037 across 18 antigen variants, outperforming both AF2-multimer and AF2-monomer models. For the SPR dataset, FAMPNN `0_3_cath` performs the best across all deep learning models and the baselines, with a correlation coefficient of 0.269 ± 0.027. We observe a notable drop in performance of models and baselines in the antibody ranking tasks (e.g 45% drop of AF2 monomer model performance on CoV-AbDab). This discrepancy likely stems from both biological and dataset-level factors. Antibody variable domains are highly flexible and structurally diverse, whereas antigenic regions like the RBD and NTD are comparatively rigid and easier to model. Importantly, antigen ranking involves mutations at a fixed epitope (the antibody’s binding site), while antibody ranking involves multiple epitopes across the antigen surface. Consequently, mutation-based predictors lose explanatory power when multiple biological factors affect binding, leading to weaker correlations.

**Table 3.** Performance metrics for antibody ranking across datasets and models. Reported values include the mean ± SEM for both ROC-AUC and Spearman correlation coefficient 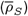. Higher positive values indicate stronger performance, where the best-performing model is highlighted in bold and second best is underlined. – indicates data are not available, where DMS baseline scores were constant across antigen lineages resulting in no correlations across both datasets.

| Model | Cov-AbDab<br>Binding ROC-AUC | SPR Data<br>$\bar{\rho}_S$ |
| --- | --- | --- |
| Epitope Alteration Count | 0.514 ± 0.040 | 0.143 ± 0.017 |
| Cao DMS Baseline | – | – |
| Bloom DMS Baseline | – | – |
| AF2 Multimer | 0.488 ± 0.042 | 0.040 ± 0.013 |
| AF2 Monomer | 0.468 ± 0.036 | 0.134 ± 0.027 |
| Boltz1 | **0.562 ± 0.037** | 0.212 ± 0.044 |
| ProteinMPNN | 0.490 ± 0.023 | 0.174 ± 0.022 |
| MPNNsol | 0.477 ± 0.028 | 0.195 ± 0.025 |
| AbMPNN | 0.496 ± 0.027 | <u>0.222 ± 0.026</u> |
| FAMPNN 0_3_cath | <u>0.535 ± 0.048</u> | **0.269 ± 0.027** |
| FAMPNN 0_3 | 0.433 ± 0.039 | 0.145 ± 0.019 |
| FAMPNN 0_0 | 0.435 ± 0.033 | 0.120 ± 0.019 |

Beyond predicting binding affinity, we explored the ability of our models to predict the neutralisation potency of individual antibodies across a broad spectrum of viral variants using the DRDB dataset (see table S2), a task that reflects a more complex and biologically downstream effect than binding alone. It is important to note that IC_50_ values in DRDB stem from diverse assay conditions, such as lentivirus and HIV-based pseudovirus systems, which introduce variability and limit direct comparability across antibodies. Encouragingly, FAMPNN checkpoints performed competitively, especially when compared to structure-based folding models, achieving a mean 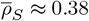 across checkpoints. Notably, a simple baseline based on epitope alteration counts achieved even stronger performance, with a correlation of 0.487 ± 0.043, highlighting both the challenge and the opportunity for further model development in this domain.

## Conclusions

CoV-UniBind was developed as a centralised, accessible and comprehensive resource for structural binding and neutralisation data of antibodies targeting SARS-CoV-2 and its variants. By curating and integrating data from a diverse array of sources, the dataset offers a rich and multifaceted view of antibody–antigen interactions. This heterogeneity enables the representation of a broad spectrum of binding affinities, neutralising potencies, and structural variations, which are often underrepresented in individual datasets. Importantly, we demonstrated that this diversity provides a strong foundation for benchmarking predictive models, enabling the evaluation of their performance across a wide range of biological and experimental contexts.

Our benchmarking results indicate that both folding and inverse folding models can show strong performance, depending on the task, suggesting that both approaches should be considered for structure-based antibody–antigen binding prediction. Performance varied across datasets, highlighting the need for task-specific fine-tuning. Despite promising results on the dataset, considerable room for improvement remains, particularly for inverse folding models. In parallel, our findings indicate that simple baseline models can provide surprisingly informative predictions. This underscores the importance of including a range of model types in benchmarking efforts, as even minimal approaches offer valuable context for evaluating and interpreting the performance of more complex methods.

With new SARS-CoV-2 variants continuing to emerge (e.g. LP.8.1 and NB.1.8.1), which carry mutations with potential implications for immune evasion and reduced vaccine efficacy (27, 28), there is a need for robust tools to anticipate and mitigate such threats. We envision CoV-UniBind not only as a benchmark for evaluating predictive models but also as a versatile platform to support the rational design of next-generation antibodies and vaccines (11, 47). By offering a comprehensive, integrated view of antibody–antigen interactions across diverse structural and functional landscapes, CoV-UniBind lays the groundwork for accelerating therapeutic development in response to SARS-CoV-2 evolution. Moreover, the framework established here can be readily adapted to future viral threats, contributing to a more agile and informed pandemic preparedness strategy.

## Competing interests

The following authors were employed by InstaDeep Ltd. at the time of this work: Aryan Bhasin, Francesco Saccon, Callum Canavan, Andrew Robson, Joao Euko, and Yunguan Fu. Alexandra C. Walls was employed by BioNTech US at the time of this work. All work presented in this publication was funded by InstaDeep Ltd. Yunguan Fu and Alexandra C. Walls are stockholders in BioNTech SE.

## Author contributions statement

Aryan Bhasin (Methodology [lead], Writing – original draft [equal], Writing—review & editing [equal], Data curation [supporting], Formal analysis [supporting]), Francesco Saccon (Data curation [lead], Writing – original draft [equal], Writing—review & editing [equal], Formal analysis [supporting]), Callum Canavan (Methodology [equal], Writing—review & editing [equal], Formal analysis [supporting]), Andrew Robson (Data curation [supporting], Writing—review & editing [equal]), Joao Euko (Software [lead], Data curation [supporting]) Alexandra C. Walls (Conceptualisation [equal], Writing—review & editing [equal]), Yunguan Fu (Conceptualisation [lead], Supervision [lead], Data curation [lead], Methodology [supporting], Writing – original draft [equal]), Writing—review & editing [equal])

## ACKNOWLEDGEMENTS

We would like to thank Alexander Muik, Bonny Gaby Lui, and Sébastien Felt for their contributions to the conceptual development of this work. We also thank Rafal Okuniewski, Michael Murray, Batu Yildirim, Elis Roberts, Isaac Rayment, Bachir Djermani, Sergio Chaves García-Mascaraque, Jonathan Faustin, Skander Marsit, Achille Soulié, and Léonie Cheminais for their support throughout the project.

## Supplementary Data

**Supplementary Table S1.** Performance of binding classification for the antigen ranking task for the Cov-AbDab Lineage-Antibody Binding dataset, split into RBD-targeting and NTD-targeting antibody subsets. Higher positive values indicate stronger performance, where the best-performing model is highlighted in bold and second best is underlined. A Mann-Whitney U-test was performed to test statistical significance; * indicates p < 0.05.

| Model | Cov-AbDab Binding<br>(RBD) ROC-AUC | Cov-AbDab Binding<br>(NTD) ROC-AUC |
| --- | --- | --- |
| Epitope Alteration Count | 0.803 ± 0.040 | **0.795 ± 0.069** |
| Cao DMS Baseline | 0.834 ± 0.041 | 0.513 ± 0.124* |
| Bloom DMS Baseline | <u>0.840 ± 0.041</u> | 0.499 ± 0.124* |
| AF2 Multimer | 0.556 ± 0.039 | 0.550 ± 0.158 |
| AF2 Monomer | **0.860 ± 0.032** | <u>0.765 ± 0.118</u> |
| Boltz1 | 0.693 ± 0.040 | 0.589 ± 0.140 |
| ProteinMPNN | 0.736 ± 0.040 | 0.640 ± 0.129 |
| MPNNsol | 0.703 ± 0.039 | 0.640 ± 0.107 |
| AbMPNN | 0.779 ± 0.040 | 0.654 ± 0.097 |
| FAMPNN 0_3_cath | 0.778 ± 0.037 | 0.515 ± 0.130 |
| FAMPNN 0_3 | 0.668 ± 0.053 | 0.644 ± 0.115 |
| FAMPNN 0_0 | 0.756 ± 0.044 | 0.593 ± 0.147 |

**Supplementary Table S2.** Performance metrics for antigen ranking using neutralisation IC_50_ data from DRDB. Reported values include the mean ± SEM for Spearman correlation coefficients 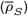. Higher positive values indicate stronger performance, where the best-performing model is highlighted in bold and second best is underlined.

| Model | DRDB Neutralisation<br>Potency $\bar{\rho}_S$ |
| --- | --- |
| Epitope Alteration Count | **0.487 ± 0.043** |
| Cao DMS Baseline | 0.391 ± 0.035 |
| Bloom DMS Baseline | 0.422 ± 0.035 |
| AF2 Multimer | 0.023 ± 0.040 |
| AF2 Monomer | 0.352 ± 0.039 |
| Boltz1 | 0.180 ± 0.040 |
| ProteinMPNN | 0.268 ± 0.050 |
| MPNNsol | 0.285 ± 0.047 |
| AbMPNN | 0.298 ± 0.045 |
| FAMPNN 0_3_cath | 0.315 ± 0.044 |
| FAMPNN 0_3 | 0.402 ± 0.038 |
| FAMPNN 0_0 | <u>0.424 ± 0.039</u> |

## Bibliography

1. Duduzile Ndwandwe and Charles S Wiysonge. Covid-19 vaccines. Current opinion in immunology, 71:111–116, 2021.

2. Richard W Titball, David I Bernstein, Nicolas VJ Fanget, Roy A Hall, Stephanie Longet, Paul A MacAry, Richard E Rupp, Marit van Gils, Veronika von Messling, David H Walker, et al. Progress with covid vaccine development and implementation. npj Vaccines, 9(1):69, 2024.

3. Yu-Chyi Hwang, Ruei-Min Lu, Shih-Chieh Su, Pao-Yin Chiang, Shih-Han Ko, Feng-Yi Ke, Kang-Hao Liang, Tzung-Yang Hsieh, and Han-Chung Wu. Monoclonal antibodies for covid-19 therapy and sars-cov-2 detection. Journal of Biomedical Science, 29(1):1, 2022.

4. Yuelong Shu and John McCauley. Gisaid: Global initiative on sharing all influenza data–from vision to reality. Eurosurveillance, 22(13):30494, 2017.

5. Matthew IJ Raybould, Aleksandr Kovaltsuk, Claire Marks, and Charlotte M Deane. Covabdab: the coronavirus antibody database. Bioinformatics, 37(5):734–735, 2021.

6. Peter W Harrison, Rodrigo Lopez, Nadim Rahman, Stefan Gutnick Allen, Raheela Aslam, Nicola Buso, Carla Cummins, Yasmin Fathy, Eloy Felix, Mihai Glont, et al. The covid-19 data portal: accelerating sars-cov-2 and covid-19 research through rapid open access data sharing. Nucleic acids research, 49(W1):W619–W623, 2021.

7. Kristie L Oxford, Jeremy D Zucker, Jeremy R Teuton, and Marek Ostaszewski. Covid19 disease map, a computational knowledge repository of sars-cov-2 virus-host interaction mechanisms. Molecular Systems Biology, 17(PNNL-SA-157722), 2021.

8. Adil R Sarhan, Thaer A Hussein, Mohammed H Flaih, and Khwam R Hussein. A biochemical analysis of patients with covid-19 infection. Biochemistry Research International, 2021 (1):1383830, 2021.

9. Rabie Adel El Arab, May Alkhunaizi, Yousef N Alhashem, Alissar Al Khatib, Munirah Bubsheet, and Salwa Hassanein. Artificial intelligence in vaccine research and development: an umbrella review. Frontiers in Immunology, 16:1567116, 2025.

10. Karim Beguir, Marcin J Skwark, Yunguan Fu, Thomas Pierrot, Nicolas Lopez Carranza, Alexandre Laterre, Ibtissem Kadri, Abir Korched, Anna U Lowegard, Bonny Gaby Lui, et al. Early computational detection of potential high-risk sars-cov-2 variants. Computers in biology and medicine, 155:106618, 2023.

11. Frédéric A Dreyer, Constantin Schneider, Aleksandr Kovaltsuk, Daniel Cutting, Matthew J Byrne, Daniel A Nissley, Henry Kenlay, Claire Marks, David Errington, Richard J Gildea, et al. Computational design of therapeutic antibodies with improved developability: efficient traversal of binder landscapes and rescue of escape mutations. In mAbs, volume 17, page 2511220. Taylor & Francis, 2025.

12. Chuan Wang, Jiangyuan Wang, Wenjun Song, Guanzheng Luo, and Taijiao Jiang. Episcan: accurate high-throughput mapping of antibody-specific epitopes using sequence information. NPJ Systems Biology and Applications, 10(1):101, 2024.

13. Hao Lv, Lei Shi, Joshua William Berkenpas, Fu-Ying Dao, Hasan Zulfiqar, Hui Ding, Yang Zhang, Liming Yang, and Renzhi Cao. Application of artificial intelligence and machine learning for covid-19 drug discovery and vaccine design. Briefings in bioinformatics, 22(6): bbab320, 2021.

14. Helen M Berman, John Westbrook, Zukang Feng, Gary Gilliland, Talapady N Bhat, Helge Weissig, Ilya N Shindyalov, and Philip E Bourne. The protein data bank. Nucleic acids research, 28(1):235–242, 2000.

15. Pascal Notin, Aaron Kollasch, Daniel Ritter, Lood Van Niekerk, Steffanie Paul, Han Spinner, Nathan Rollins, Ada Shaw, Rose Orenbuch, Ruben Weitzman, et al. Proteingym: Largescale benchmarks for protein fitness prediction and design. Advances in Neural Information Processing Systems, 36:64331–64379, 2023.

16. Johnathan D Guest, Thom Vreven, Jing Zhou, Iain Moal, Jeliazko R Jeliazkov, Jeffrey J Gray, Zhiping Weng, and Brian G Pierce. An expanded benchmark for antibody-antigen docking and affinity prediction reveals insights into antibody recognition determinants. Structure, 29(6):606–621, 2021.

17. Jarjapu Mahita, Brendan Ha, Anais Gambiez, Sharon L Schendel, Haoyang Li, Kathryn M Hastie, S Moses Dennison, Kan Li, Natalia Kuzmina, Sivakumar Periasamy, et al. Coronavirus immunotherapeutic consortium database. Database, 2023:baac112, 2023.

18. Anna Huhn, Daniel Nissley, Daniel B Wilson, Mikhail A Kutuzov, Robert Donat, Tiong Kit Tan, Ying Zhang, Michael I Barton, Chang Liu, Wanwisa Dejnirattisai, et al. The molecular reach of antibodies crucially underpins their viral neutralisation capacity. Nature Communications, 16(1):338, 2025.

19. Allison J Greaney, Tyler N Starr, and Jesse D Bloom. An antibody-escape estimator for mutations to the sars-cov-2 receptor-binding domain. Virus evolution, 8(1):veac021, 2022.

20. Philip L Tzou, Kaiming Tao, Sergei L Kosakovsky Pond, and Robert W Shafer. Coronavirus resistance database (cov-rdb): Sars-cov-2 susceptibility to monoclonal antibodies, convalescent plasma, and plasma from vaccinated persons. Plos one, 17(3):e0261045, 2022.

21. James Dunbar, Konrad Krawczyk, Jinwoo Leem, Terry Baker, Angelika Fuchs, Guy Georges, Jiye Shi, and Charlotte M Deane. Sabdab: the structural antibody database. Nucleic acids research, 42(D1):D1140–D1146, 2014.

22. John Jumper, Richard Evans, Alexander Pritzel, Tim Green, Michael Figurnov, Olaf Ronneberger, Kathryn Tunyasuvunakool, Russ Bates, Augustin Žídek, Anna Potapenko, et al. Highly accurate protein structure prediction with alphafold. Nature, 596(7873):583–589, 2021.

23. Jeremy Wohlwend, Gabriele Corso, Saro Passaro, Mateo Reveiz, Ken Leidal, Wojtek Swiderski, Tally Portnoi, Itamar Chinn, Jacob Silterra, Tommi Jaakkola, et al. Boltz-1: Democratizing biomolecular interaction modeling. bioRxiv, pages 2024–11, 2024.

24. Justas Dauparas, Ivan Anishchenko, Nathaniel Bennett, Hua Bai, Robert J Ragotte, Lukas F Milles, Basile IM Wicky, Alexis Courbet, Rob J de Haas, Neville Bethel, et al. Robust deep learning–based protein sequence design using proteinmpnn. Science, 378(6615):49–56, 2022.

25. Casper A. Goverde, Martin Pacesa, Nicolas Goldbach, Lars J. Dornfeld, Petra E. M. Balbi, Sandrine Georgeon, Stéphane Rosset, Srajan Kapoor, Jagrity Choudhury, Justas Dauparas, Christian Schellhaas, Simon Kozlov, David Baker, Sergey Ovchinnikov, Alex J. Vecchio, and Bruno E. Correia. Computational design of soluble and functional membrane protein analogues. Nature, 631(8020):449–458, July 2024. ISSN 1476-4687. doi: 10.1038/s41586-024-07601-y.

26. Frédéric A. Dreyer, Daniel Cutting, Constantin Schneider, Henry Kenlay, and Charlotte M. Deane. Inverse folding for antibody sequence design using deep learning. 2023 ICML Workshop on Computational Biology, 2023.

27. Jingyi Liu, Yuanling Yu, Sijie Yang, Fanchong Jian, Weiliang Song, Lingling Yu, Fei Shao, and Yunlong Cao. Virological and antigenic characteristics of sars-cov-2 variants lf. 7.2. 1, np. 1, and lp. 8.1. The Lancet Infectious Diseases, 2025.

28. Caiwan Guo, Yuanling Yu, Jingyi Liu, Fanchong Jian, Sijie Yang, Weiliang Song, Lingling Yu, Fei Shao, and Yunlong Cao. Antigenic and virological characteristics of sars-cov-2 variants ba. 3.2, xfg, and nb. 1.8. 1. The Lancet Infectious Diseases, 2025.

29. Stephen J Goodswen, Paul J Kennedy, and John T Ellis. A guide to current methodology and usage of reverse vaccinology towards in silico vaccine discovery. FEMS Microbiology Reviews, 47(2):fuad004, 2023.

30. Karthik Gangavarapu, Alaa Abdel Latif, Julia L Mullen, Manar Alkuzweny, Emory Hufbauer, Ginger Tsueng, Emily Haag, Mark Zeller, Christine M Aceves, Karina Zaiets, et al. Out-break. info genomic reports: scalable and dynamic surveillance of sars-cov-2 variants and mutations. Nature Methods, 20(4):512–522, 2023.

31. Yunlong Cao, Fanchong Jian, Jing Wang, Yuanling Yu, Weiliang Song, Ayijiang Yisimayi, Jing Wang, Ran An, Xiaosu Chen, Na Zhang, et al. Imprinted sars-cov-2 humoral immunity induces convergent omicron rbd evolution. Nature, 614(7948):521–529, 2023.

32. Qi Zhang, Bin Ju, Jiwan Ge, Jasper Fuk-Woo Chan, Lin Cheng, Ruoke Wang, Weijin Huang, Mengqi Fang, Peng Chen, Bing Zhou, et al. Potent and protective ighv3-53/3-66 public antibodies and their shared escape mutant on the spike of sars-cov-2. Nature communications, 12(1):4210, 2021.

33. Bin Ju, Qi Zhang, Jiwan Ge, Ruoke Wang, Jing Sun, Xiangyang Ge, Jiazhen Yu, Sisi Shan, Bing Zhou, Shuo Song, et al. Human neutralizing antibodies elicited by sars-cov-2 infection. Nature, 584(7819):115–119, 2020.

34. Fanchong Jian, Jing Wang, Ayijiang Yisimayi, Weiliang Song, Yanli Xu, Xiaosu Chen, Xiao Niu, Sijie Yang, Yuanling Yu, Peng Wang, et al. Evolving antibody response to sars-cov-2 antigenic shift from xbb to jn. 1. Nature, 637(8047):921–929, 2025.

35. Bissan Al-Lazikani, Arthur M Lesk, and Cyrus Chothia. Standard conformations for the canonical structures of immunoglobulins. Journal of molecular biology, 273(4):927–948, 1997.

36. Peter W Rose, Bojan Beran, Chunxiao Bi, Wolfgang F Bluhm, Dimitris Dimitropoulos, David S Goodsell, Andreas Prlic, Martha Quesada, Gregory B Quinn, John D Westbrook, et al. The rcsb protein data bank: redesigned web site and web services. Nucleic acids research, 39(suppl_1):D392–D401, 2010.

37. James Dunbar and Charlotte M Deane. Anarci: antigen receptor numbering and receptor classification. Bioinformatics, 32(2):298–300, 2016.

38. Robert M MacCallum, Andrew CR Martin, and Janet M Thornton. Antibody-antigen interactions: contact analysis and binding site topography. Journal of molecular biology, 262(5): 732–745, 1996.

39. Jiansheng Jiang, Christopher T Boughter, Javeed Ahmad, Kannan Natarajan, Lisa F Boyd, Martin Meier-Schellersheim, and David H Margulies. Sars-cov-2 antibodies recognize 23 distinct epitopic sites on the receptor binding domain. Communications Biology, 6(1):953, 2023.

40. Min Huang, Lili Wu, Anqi Zheng, Yufeng Xie, Qingwen He, Xiaoyu Rong, Pu Han, Pei Du, Pengcheng Han, Zengyuan Zhang, et al. Atlas of currently available human neutralizing antibodies against sars-cov-2 and escape by omicron sub-variants ba. 1/ba. 1.1/ba. 2/ba. 3. Immunity, 55(8):1501–1514, 2022.

41. Qingwen He, Lili Wu, Zepeng Xu, Xiaoyun Wang, Yufeng Xie, Yan Chai, Anqi Zheng, Jianjie Zhou, Shitong Qiao, Min Huang, et al. An updated atlas of antibody evasion by sars-cov-2 omicron sub-variants including bq. 1.1 and xbb. Cell Reports Medicine, 4(4), 2023.

42. Patrick Bryant, Gabriele Pozzati, and Arne Elofsson. Improved prediction of protein-protein interactions using alphafold2. Nature Communications, 13(1):1265, 2022. doi: 10.1038/s41467-022-28865-w.

43. Jeremy Wohlwend, Gabriele Corso, Saro Passaro, Noah Getz, Mateo Reveiz, Ken Leidal, Wojtek Swiderski, Liam Atkinson, Tally Portnoi, Itamar Chinn, Jacob Silterra, Tommi Jaakkola, and Regina Barzilay. Boltz-1 democratizing biomolecular interaction modeling. bioRxiv, 2025. doi: 10.1101/2024.11.19.624167.

44. Justin Gilmer, Samuel S Schoenholz, Patrick F Riley, Oriol Vinyals, and George E Dahl. Neural message passing for quantum chemistry. In International conference on machine learning, pages 1263–1272. PMLR, 2017.

45. Richard W. Shuai, Talal Widatalla, Po-Ssu Huang, and Brian L. Hie. Sidechain conditioning and modeling for full-atom protein sequence design with fampnn. bioRxiv, 2025. doi: 10.1101/2025.02.13.637498.

46. L América Chi, Jonathan E Barnes, Jagdish Suresh Patel, and F Marty Ytreberg. Exploring the ability of the md+ foldx method to predict sars-cov-2 antibody escape mutations using large-scale data. Scientific Reports, 14(1):23122, 2024.

47. James A Williams, Marco Biancucci, Laura Lessen, Sai Tian, Ankita Balsaraf, Lynn Chen, Chelsy Chesterman, Giulietta Maruggi, Sarah Vandepaer, Ying Huang, et al. Structural and computational design of a sars-cov-2 spike antigen with improved expression and immunogenicity. Science Advances, 9(23):eadg0330, 2023.

